# A Ras-like GTPase in African trypanosomes

**DOI:** 10.1101/2025.03.03.640364

**Authors:** Margaret Lowenstein, Alejandro Ramirez, Mark C. Field

**Affiliations:** Department of Pathology, University of Cambridge, Tennis Court Road, Cambridge, UK; School of Life Sciences, University of Dundee, Dundee, UK; Institute of Parasitology, Biology Centre, Czech Academy of Sciences, České Budějovice, Czech Republic

**Keywords:** small G protein, trypanosomes, signalling, evolution

## Abstract

Small Ras-like GTPases are conserved across eukaryotes and mediate many functions. *Trypanosoma brucei* encodes only three putative open reading frames belonging to Ras and Rho subfamilies, all highly divergent from other taxa. We investigated the most Ras-like gene product in *T. brucei*, TbRlp, evaluating essentiality, location and impact of blocking the GDP/GTP cycle. TbRlp is expressed in both life stages, with a clear fitness contribution in bloodstream forms in *in vitro* culture, without a specific block to the cell cycle or clear morphological defect. Overexpression of TbRlp and mutant forms results in moderate disruption of the cell cycle and altered proliferation indicating that GTP/GDP cycling and expression levels of these forms of TbRlp are important to proliferation. Epitope-tagged TbRlp is located predominantly on structures in the posterior of the cell, and suggesting the late endosomal system. Analyses also indicates a link between cell density and levels of TbRlp in the BSF, with a decrease in TbRlp protein but not RNA upon increased cell density. These data suggest that TbRlp is important in control of proliferation and opens possibilities for future research.

## Main article

*Trypanosoma brucei spp*. proceeds through multiple life-cycle stages during progress between mammalian and insect hosts, alternating between replicating and quiescent forms, including forms dwelling in adipose tissue, skin and bloodstream (Trindade et al., 2016). Life cycle progression requires extensive cellular remodelling, encompassing the surface coat, primary metabolism and other processes. The Ras superfamily of GTPases are important mediators of signal transduction (Macara *et al*., 1996); trypanosomes possess a small GTPase complement (Field 2005), and TbRlp (Tb927.11.6030) is the closest Ras homolog (Sowa et al., 1999, Field and O’Reilly 2008). Many observations indicate that trypanosomes have unique signaling pathways, included differentiation pathways depending on trypanosome-specific machinery (Salmon 2018, Matthews 2021, Cayla et al., 2024)

In animals Ras-signaling *via* serine/threonine Raf and Vps34 PI-3 kinases suggests tight integration with protein and lipid signal cascades, and the dominant role of Ras in cell proliferation *via* mitogen-activated protein kinase (MAPK) pathways and others is well known (Singh et al., 2020). Proteins with Raf-like kinase domains are encoded in the trypanosome genome, but divergent overall protein architecture suggests that molecules coupling trypanosome Ras-like proteins to MAPK pathways remain unknown. The presence of multiple MAPK paralogs suggests that part of the pathway is conserved across evolution, while an orthologue of Vps34 is present in trypanosomes and required for several cellular functions, including Golgi complex replication (Hall *et al*., 2006). Further, cAMP-mediated pathways are central to differentiation in trypanosomes (Salmon 2018, Bachmaier et al., 2023), suggesting some similarity to Ras signalling via cAMP in yeasts (Bonomelli et al., 2023). Here we report the location, essentiality and phenotypic impact of disruption of TbRlp on trypanosomes.

The Tb927.11.6030 ORF encodes a protein of 226 amino acids with a molecular weight of 25.0 kDa. Orthologs are present in many kinetoplastida (https://tritrypdb.org/tritrypdb/app/record/gene/Tb927.11.6030#Orthologs). All canonical GTP-binding and C-terminal prenylation motifs are conserved (Figure S1). The C-terminus has a basic amino acid region preceding the prenylation signal, typical of Ras-like GTPases. Phylogenetic reconstruction using MrBayes (Huelsenbeck and Ronquist 2001) demonstrates that TbRlp is a rather divergent member of the Ras-like clade, while Alphafold3 modelling (Abramson et al., 2024) predicts a canonical small GTPase fold.

We confirmed earlier data that TbRlp is expressed in both major life stages using qRT-PCR (Sowa *et al*., 1999). We detected a low abundance PCR amplicon from mRNA from both procyclic culture form (PCF) and bloodstream form (BSF), some seven orders of magnitude less than ß-tubulin and in agreement with microarray data (Figure 1) (Koumandou *et al*., 2008).

**Figure 1:**
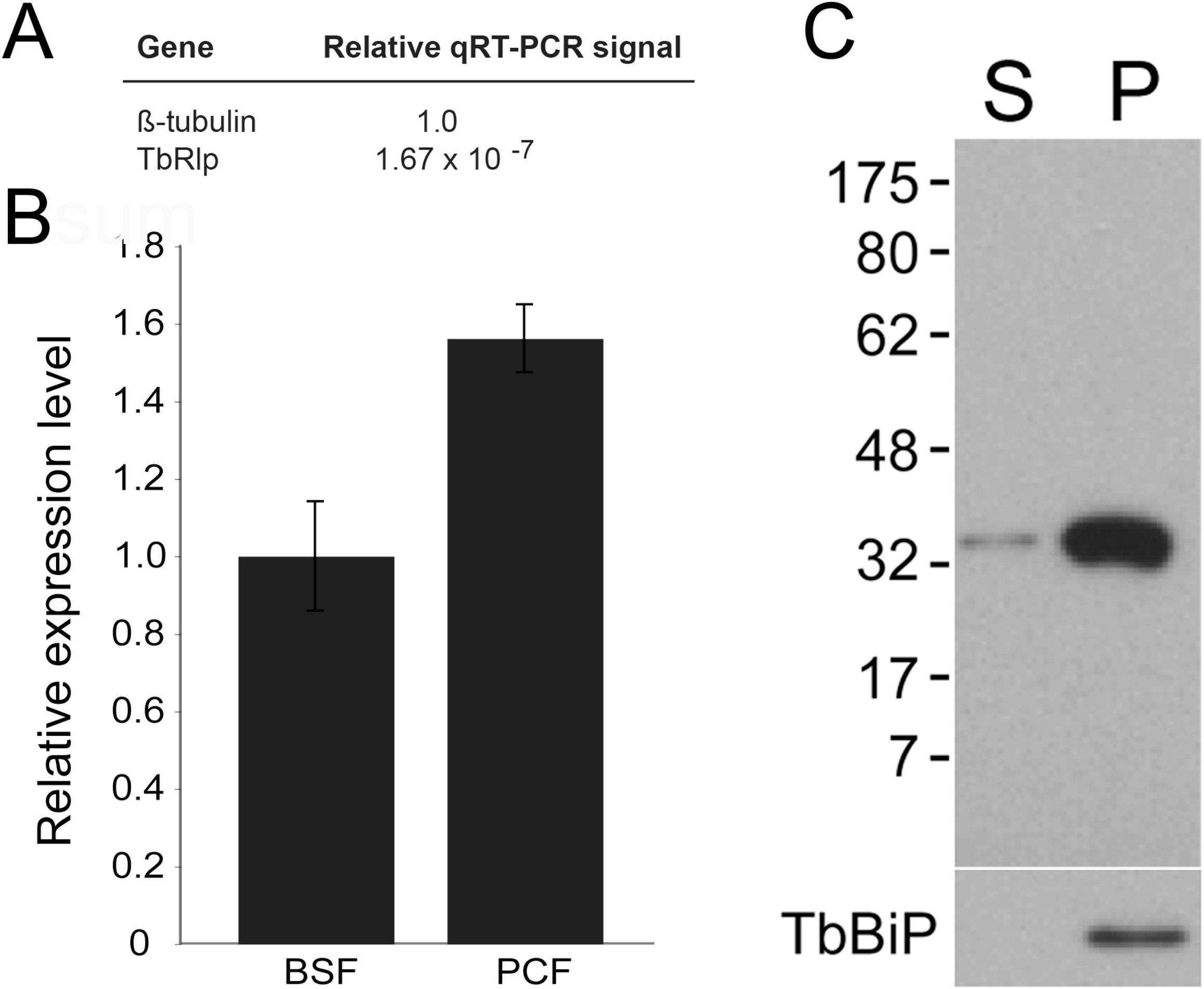
TbRlp is expressed in both major proliferative stages of *T. brucei* and required for robust growth. Panel A; qRT-PCR analysis of TbRlp mRNA levels compared to ß-tubulin. Panel B: mRNA levels in BSF and PCF stages. Data are representative of two determinations, with three technical replicates, in each case. Panel C: TbRlp is associated with the insoluble fraction. Cells expressing HA::TbRlp were lysed and pelleted at 40 000g, with equal cell equivalents of supernatant (S) and pellet (P) loaded into each lane, and probed with anti-HA and anti-TbBip. The vast majority of TbRlp is associated with the pellet. Numbers at left are molecular weight markers in kDa.

We focused most subsequent analysis on the blood stage. BSFs were transfected with the p2T7-TbRlp RNAi plasmid and transformants selected (Figure 2). By day three post-induction proliferation was reduced by ∼50%. Analysis by qRT-PCR at 24 and 48 hours post-induction indicated decreased TbRlp mRNA of ∼60%, suggesting that BSFs are highly sensitive to perturbation of TbRlp levels, and consistent with high throughput analysis (Alsford et al., 2012). Analysis of progress through the cell cycle indicated no significant alteration in distribution of cells at normal stages of replication (Figure 2). We also produced a rabbit polyclonal antibody against TbRlp as a GST fusion protein. The antisera detected a protein of the correct size in lysates prepared from induced *E. coli* (Figure S2), but also recognised a high abundance protein of ∼50KDa (Figure 3), precluding utility in localisation studies.

**Figure 2:**
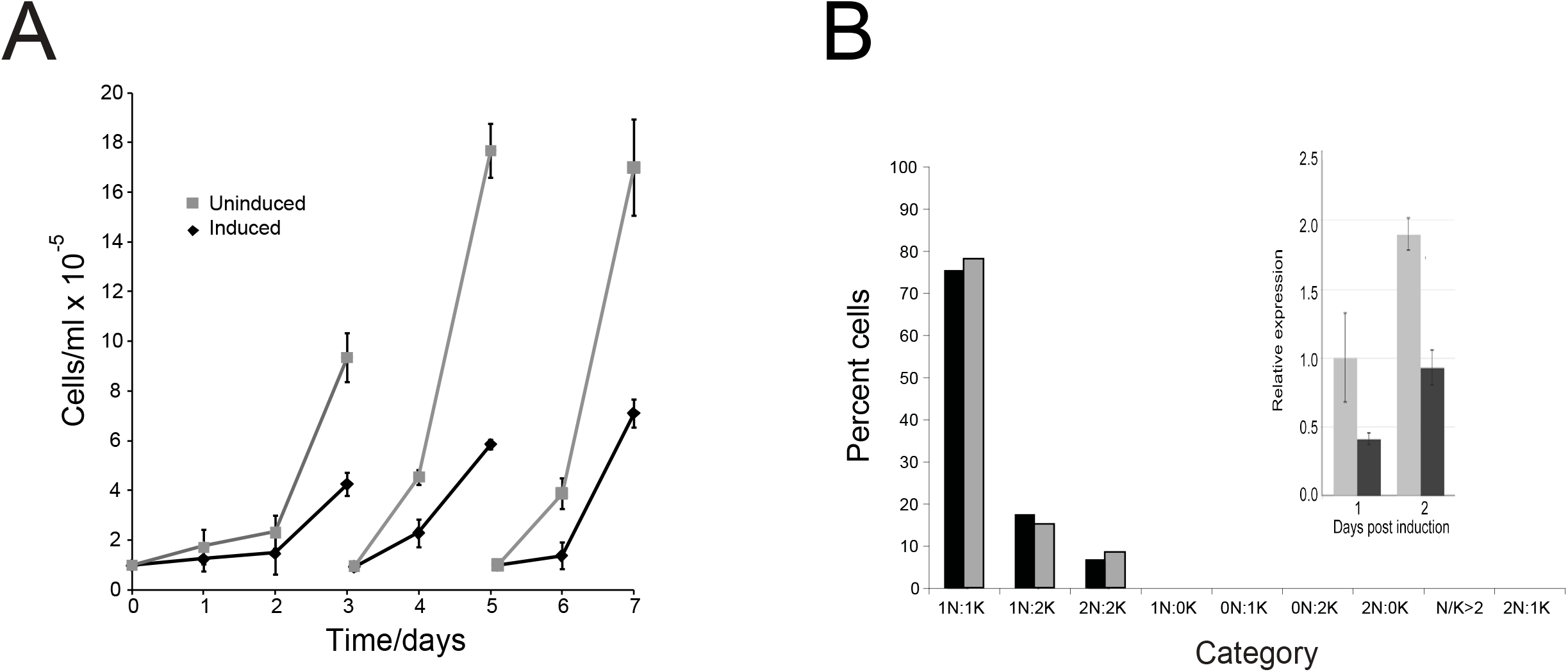
TbRlp is required for robust proliferation. Panel A: Growth curve for BSF cells harboring a TbRlp knockdown construct. Uninduced data are plotted in gray with square symbols and induced in black with diamond symbols. Data are the mean of triplicate cultures with error bars showing the standard deviation. Panel B Cell cycle analysis for knockdown cells. Data from uninduced cells are plotted in gray and induced cells in black. Over 400 cells were counted for induced and uninduced cultures. Inset RT-qPCR assay for TbRlp mRNA levels with data from uninduced cells are plotted in gray and induced cells in black. Data are representative of two determinations, with three technical replicates, in each case.

**Figure 3:**
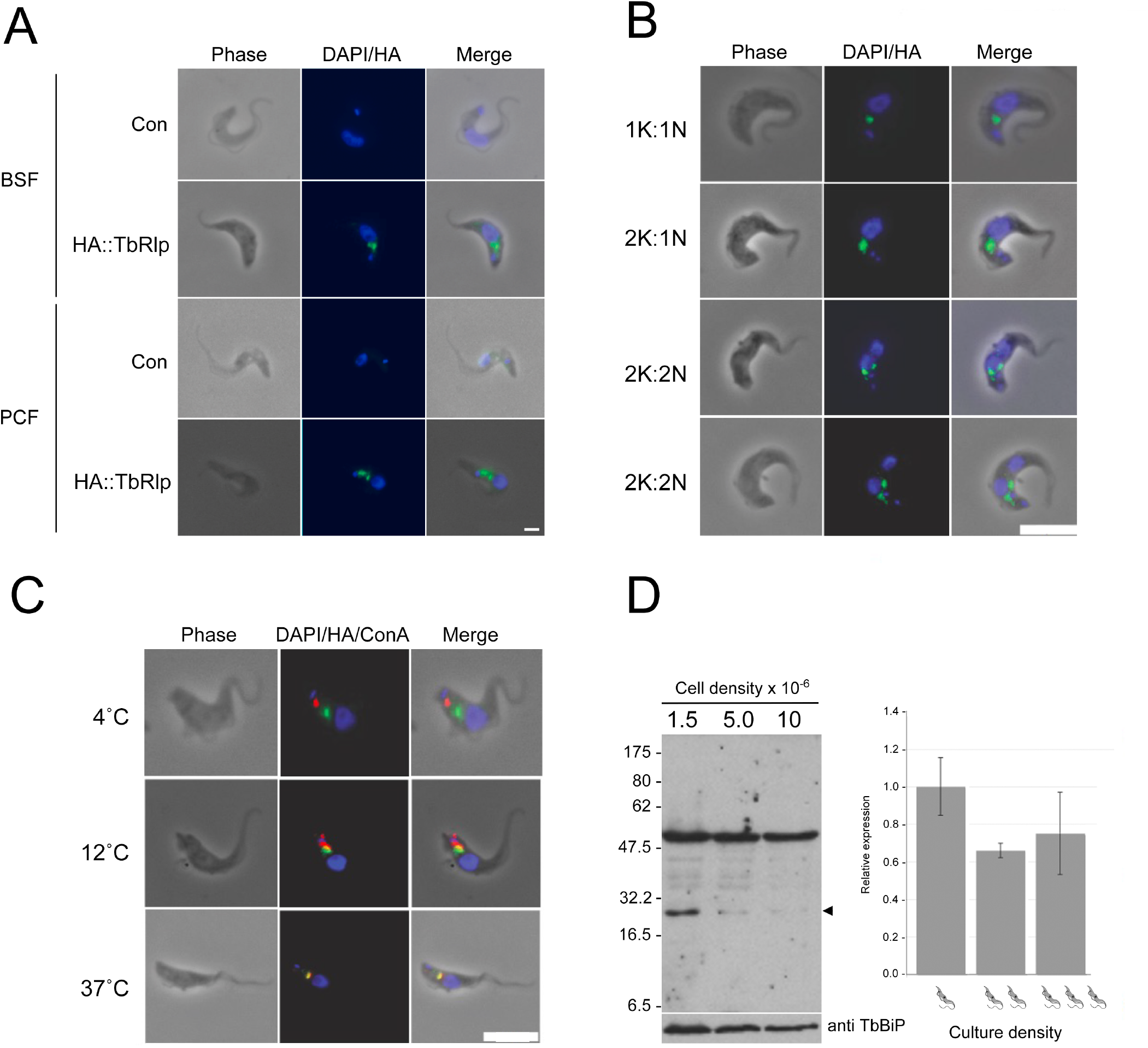
Localisation of TbRlp is likely endosomal. Panel A: BSF and PCF cells expressing HA::TbRlp (green) and demonstrating localisation of the antigen to the posterior region of the cell. DNA, stained with DAPI, is shown in blue. Scale bar is 2µm. Panel B: Analysis of cells through the cell cycle for TbRlp (green) and DNA stained with DAPI in blue. Cell cycle position is given at left. Scale bar is 6µm. Panel C: Concanavalin A uptake at various temperatures to label specific endosomal compartments. Scale bar is 6µm. Panel D: Density dependant expression of TbRlp protein. Left: Polyclonal antibody against TbRlp was used to probe whole cell lysates of BSF cells cultured at the densities shown. Arrowhead indicates the migration position of TbRlp, while a cross-reacting band is seen at ∼50kDa. anti-TbBiP was used as a loading control. 1×10^7^ cells were loaded in each lane. Right: qRT-PCR for TbRlp mRNA in BSF cells grown at various densities (low 0.3×10^6^ml, medium 0.8×10^6^ml, and high 3×10^6^ml), relative to TbRlp at low density. Error bars represent the standard deviation between three independent determinations.

We created cells expressing HA-tagged TbRlp wild-type or GTP/GDP-locked QL/ SN mutants respectively (Figure S2). HA::TbRlp is retained almost exclusively in the insoluble 40 000g pellet fraction, in common with the ER chaperone TbBiP, and consistent with presence at the membrane or other insoluble complex, such as the cytoskeleton (Figure 1). We localised the protein to the posterior region of the cell in both BSF and PCF (Figure 3), essentially unaltered in TbRlp^QL^, and only partially perturbed for the TbRlp^SN^ mutant, a form frequently partially released from membrane-association (Figure S3). During replication the TbRlp-stained structure duplicated after the kinetoplast and nucleus (Figure 3). We also examined the location of TbRlp with markers for the endomembrane system by costaining with polyclonal antibodies for the Golgi complex (GRASP), endosomes (clathrin-heavy chain) and the lysosome/vacuole (p67) (Figure S3). We found partial overlap between clathrin and TbRlp, and also p67. There was no evidence for coincidence between TbRlp and GRASP. To validate the possibililty that TbRlp is located at endosomal structures we costained cells with TbRlp that wereexposed to fluorescent Concanavalin A. When incubated at different temperatures ConA accumulates at the flagellar pocket (4°C), early/intermediate endosomes (12°C) and lysosome (37°C). We again observed considerable colocalisation between TbRlp and the lysosome, consistent with part colocalisation with p67, and suggesting a late endosome/prelysosomal location (Figure 3).

Overexpression of HA::TbRlp wildtype, HA::TbRlp^QL^ and HA::TbRlp^SN^ all led to decreased proliferation (Figure 4), with HA::TbRlp^QL^ most pronounced, suggesting that increased levels and disruption of the TbRlp GTPase cycle is detrimental to fitness. When we analysed the effect that TbRlp overexpression had on cell cycle position, we observed a small decrease in cells in interphase, i.e. 1N:1K cells, with corresponding increase in 1N:2K and 1K:0N cells. These observations suggest that the decreased proliferation is likely due to mitosis and/or nuclear division.

**Figure 4:**
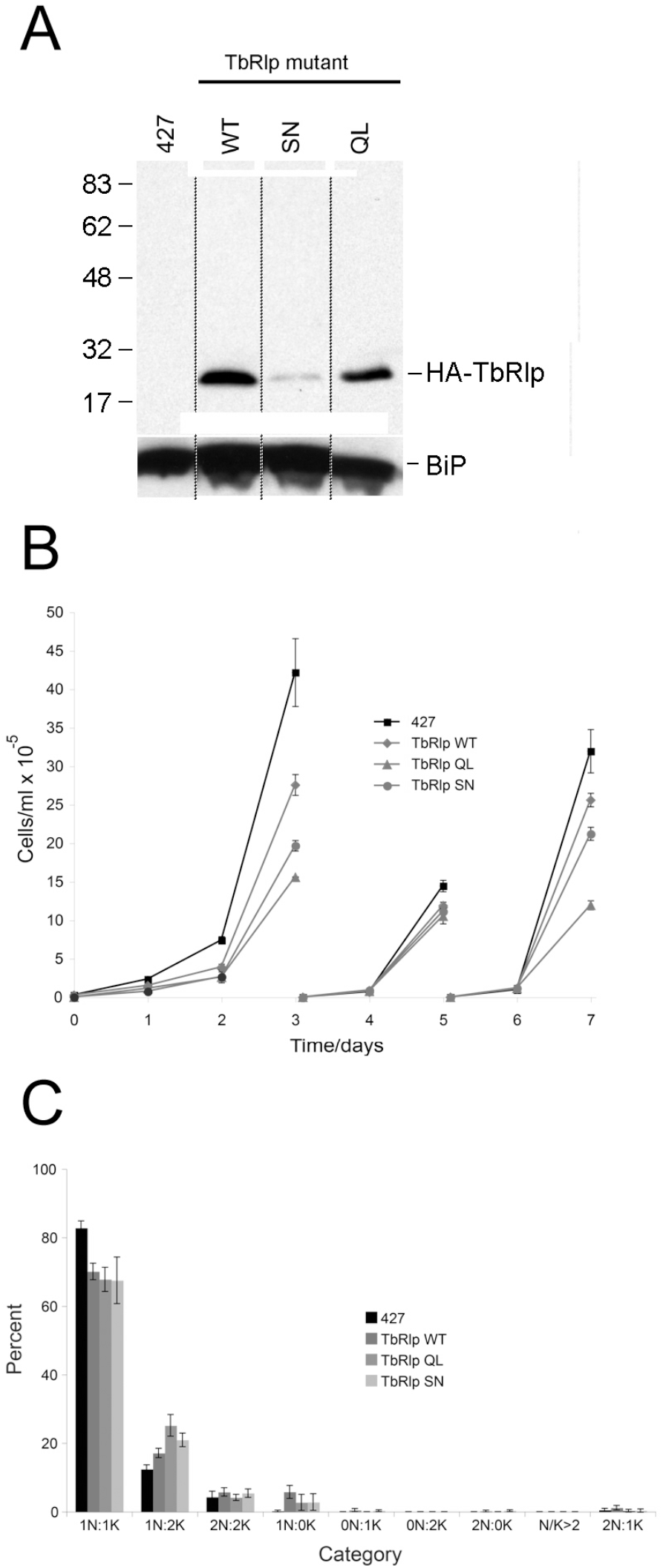
Overexpression of TbRlp and TbRlp mutants leads to proliferation and cell cycle defects in BSF trypanosomes. Panel A: Growth curves for TbRlp overexpressors over a seven day period. Curves are annotated as indicated on the figure. Panel B: Cell cycle analysis of at least 200 cells from each overexpressor class. TbRlp overexpressors were fixed, stained with DAPI and mounted. The numbers of nuclei (N) and kinetopasts (K) in each cell were determined. Black bars denote cells from control cultures and gray bars from cultures overexpressing TbRlp isoforms.

Finally, we examined the impact of cell density on TbRlp expression level, as density is a known influence on parasite differentiation. Previously we reported that mRNA expression levels driven by the rRNA promotor are density dependent, decreasing at higher densities, and also induced with conditioned medium (Ali and Field 2013). In the present case we examined expression of endogenous TbRlp using a polyclonal antibody. We observed clear evidence for density-dependant expression (Figure 3). This was apparently not a result of altered mRNA levels or could be induced using conditioned media (data not shown) and hence is likely to be mediated through protein turnover or translatability.

We note that TrypTag (http://tryptag.org) has inidicated that TbRlp is associated with the hook complex, which interacts with the flagellar pocket collar and the microtubule quartet (Albiseti et al., 2021). The data however suggest that both N- and C-terminally tagged forms of TbRlp localise here, which is highly unlikely as C-terminal tagging would disrupt prenylation and hence membrane targeting. Careful review of *bona fide* hook complex components, including TbSmee1 and TbCen2 suggest that the location reported here, and in TrypTag, is unlikely to be the hook complex, while the replication of the TbRlp punctum after kinetoplast and nuclear replication is also inconsistent with being a hook complex component. Hence we conclude that TbRlp is probably a late endosome-associated GTPase with roles in density-dependant signalling.

## Acknowledgements

This work was supported by the Wellcome Trust (to MCF), with studentship awards from the Bill and Melinda Gates Foundation (to AR and ML). We gratefully acknowledge the kind gifts of antibodies to TbBip and p67 from James Bangs and to GRASP from Graham Warren. The authors have no financial or ethical issues to declare.

## Supplementary data for

### Supplementary figure legends

**Figure S1:**
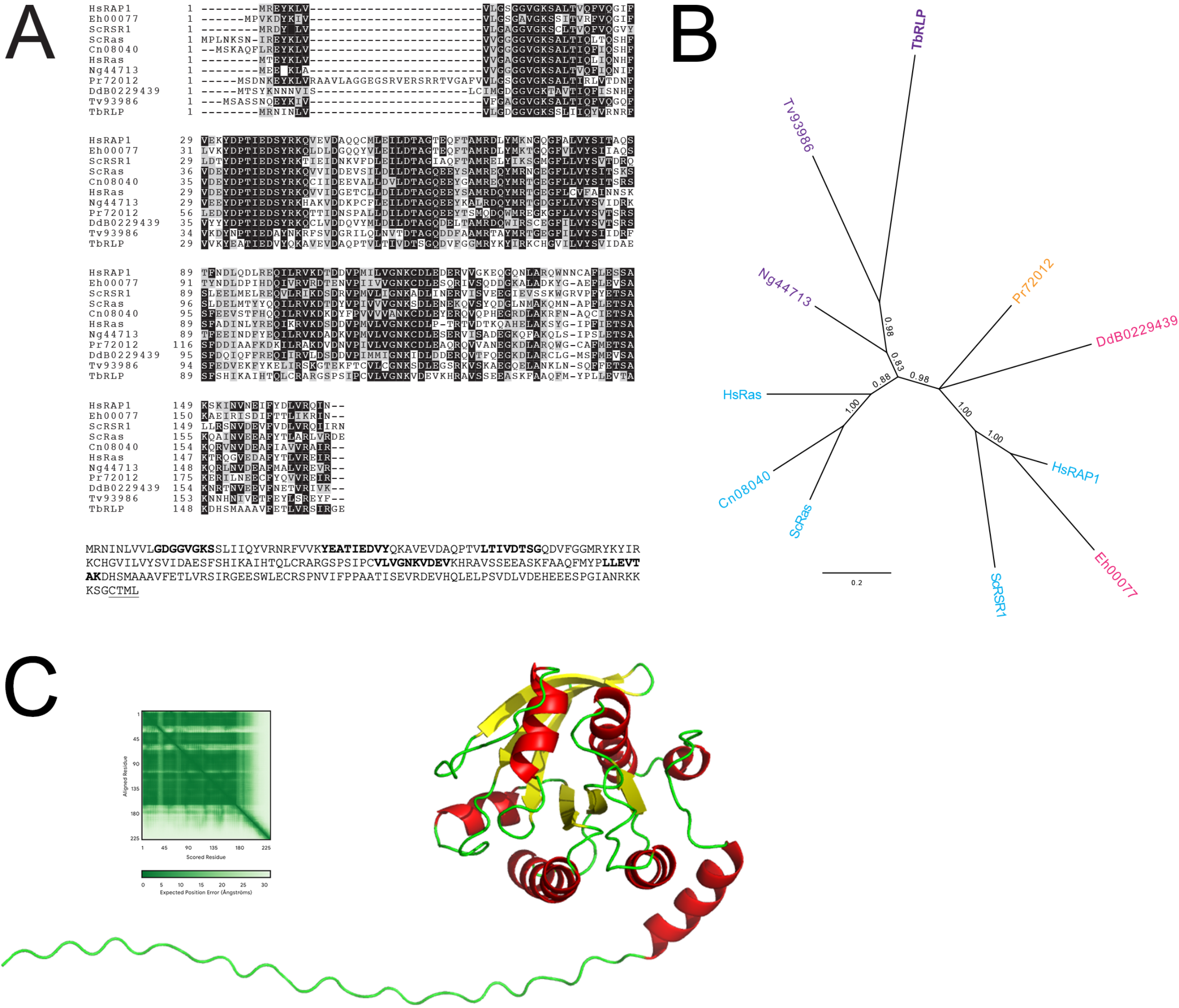
Sequence of TbRLP and some relatives. Panel A: ClustalX alignment of N-terminal portions of Ras GTPase proteins. Residues are colored with a black background if >50% of taxa retain an identical residue and gray if similar. Numbers indicate residue numbers. Sequences are annotated with a two-letter species identifier followed by either the annotated gene product name or accession number: Hs; *Homo sapiens*, Sc; *Saccharomyces cerevisiae*, Cn; *Cryptococcus neoformans*, Eh; *Entamoeba histolytica*, Dd; *Dictyostellium discoidium*, Ng; *Naegleria gruberii*, Pr; *Phytophra ramorum*, Tv; *Trichomonas vaginalis* and Tb; *Trypanosoma brucei*. Note the high level of general sequence conservation, but significant divergence within TbRlp, especially in the ILDTAG region. Lower panel: Sequences for TbRlp G1 (phosphate binding), G2 (GAP-binding), G3 (phosphate binding), G4 (guanine-binding), G5 (guanine-binding) highlighted in **bold**, and the C-terminal prenylation signal is underlined. Panel B: Unrooted MrBayes phylogenetic tree. Node numbers indicate posterior probabilities. Taxon labels are as in panel A, but colorized to indicate supergroup affiliation: Pale blue; Opistokhonta, cyan; Amoebozoa, orange; Chromalvelolata and Purple; Excavata. TbRlp is most similar to *T. vaginalis*. Panel C: Alphafold 3 prediction for TbRlp demonstrating canonical GTPase fold. α-helices are in red, β-sheets in yellow and disordered regions in green. Inset is the error calculations from the Alphafold 3 calculation.

**Figure S2:**
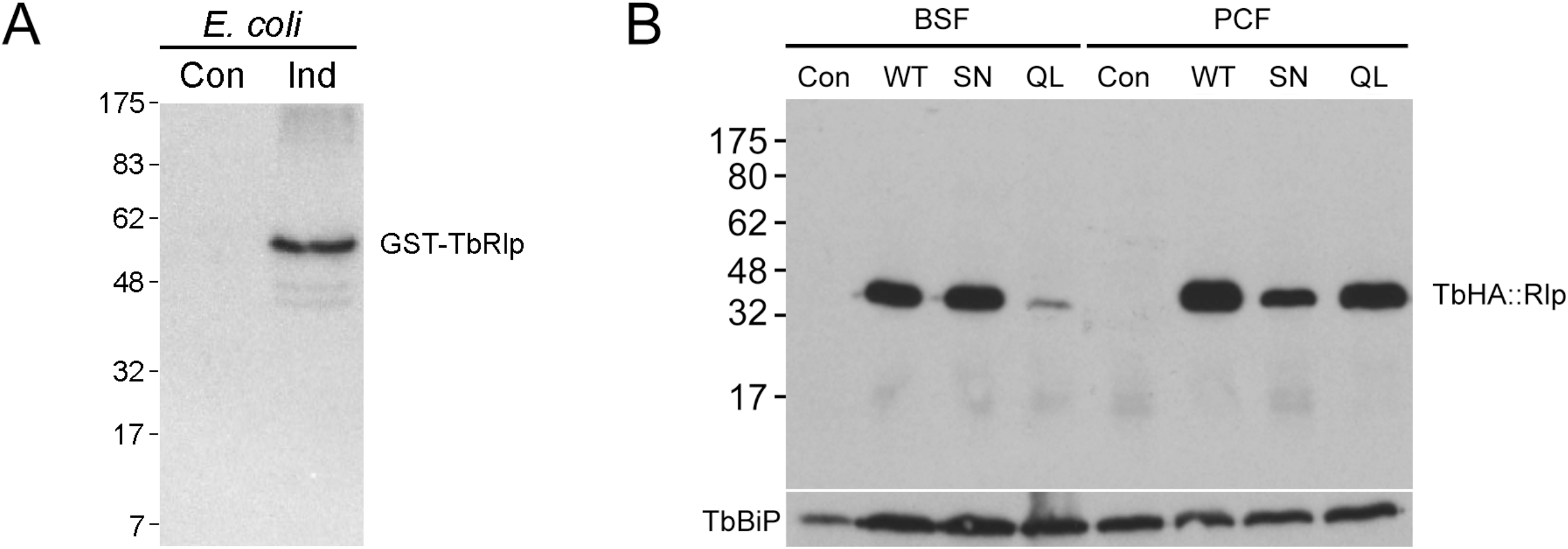
Immunoreactivity to anti-TbRlp antisera and expression of ectopically expressed TbRlp mutants. Panel A: Antibodies were affinity purified from rabbits immunized with GST-TbRlp, and used at a dilution of 1:100; bound antibodies were detected with goat anti-rabbit polyclonal antibody horseradish peroxidase conjugate developed with enhanced chemiluminescence. Lanes contain lysates from bacteria containing pGEX3X-TbRlp, either uninduced (Con) or induced (Ind) with 1mM IPTG. The expected migration position of GST-TbRlp is indicated at right, and coelectrophoresed molecular weight markers are shown at left, with molecular weight in kDa. Panel B: Western blot analysis of lysates prepared from BSF cells transfected with various TbRlp isoforms (WT; wild type, SN; serine to asparagine mutant, and QL; glutamate to leucine mutant) and wild type 427 trypanosomes. Note all data comes from a single blot; discontinuities have been introduced in rearrangement of the lanes for presentation purposes only. Top panel shows reactivity against HA and lower panel TbBiP immunoreactivity. In all cases HA immunoreactivity at ∼40kDa is detected, consistent with TbRlp harbouring three copies of the HA epitope at the N-terminus and a known slow migration of prenylated GTPases on SDS-PAGE. Migration positions of molecular weight markers are shown at left in kDa.

**Figure S3:**
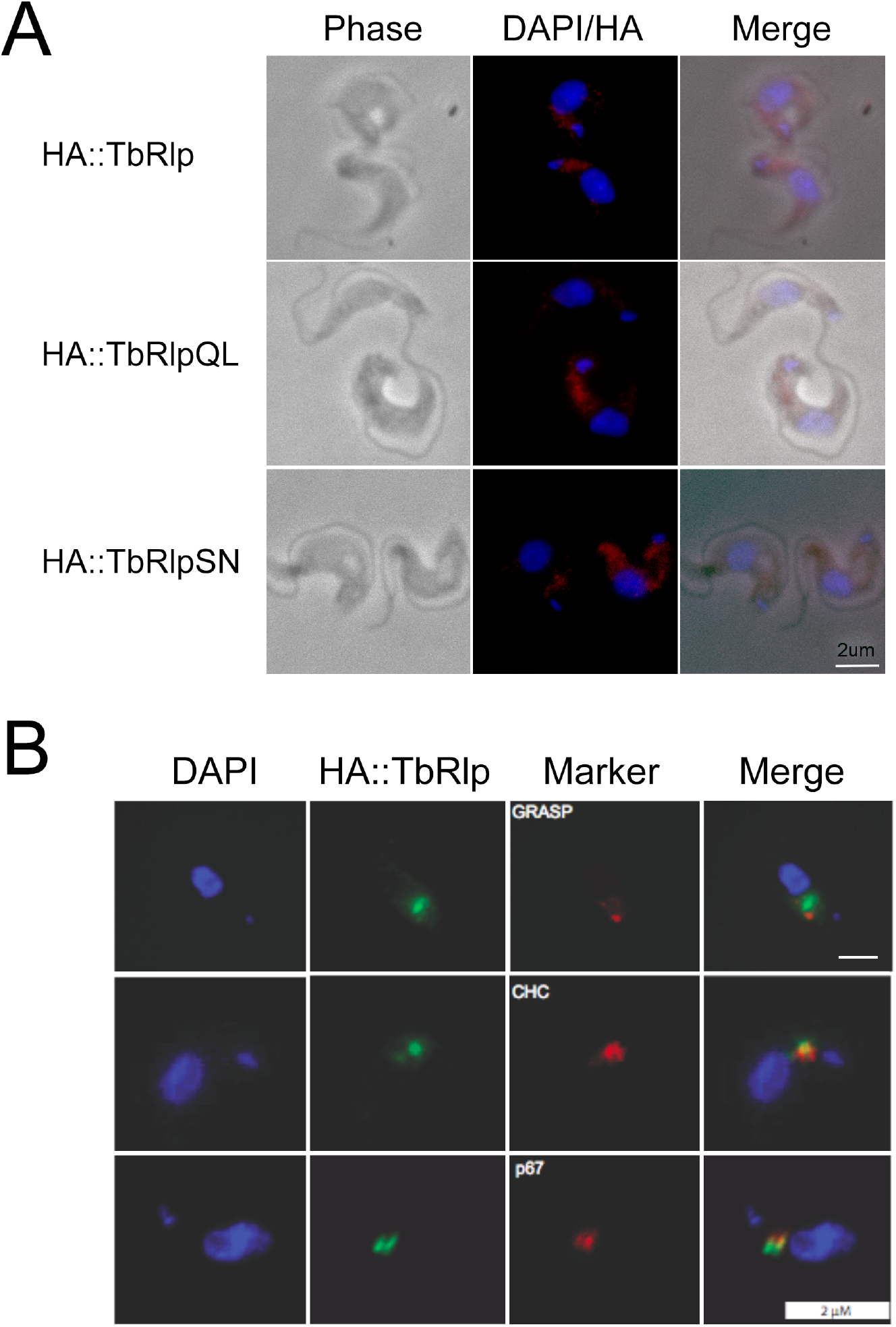
Localisation of TbRlp. Panel A: Overexpressed wild type and mutant forms of HA::TbRlp are targeted to posterior end structures. Cells were stained with anti-HA (red) to visualise the GTPase and contained with DAPI for DNA. Scale bar is 2µm. Panel B: Colocalisation of HA::TbRlp (green) and endomembrane markers; GRASP for the Golgi complex, clathin heavy chain (CHC) for endosomes and p67 for lysosome. DAPI stain for DNA is in blue, and scale bar is 2µm.

